# ONCOchannelome: A computational framework for investigating altered ion channels across tumor types

**DOI:** 10.1101/2025.11.27.685767

**Authors:** K. T. Shreya Parthasarathi, Soundharya Ramu, Mohit Kumar Jolly, Jyoti Sharma

**Affiliations:** Manipal Academy of Higher Education (MAHE), Manipal, Karnataka 576104, India; Institute of Bioinformatics, International Technology Park, Bangalore 560066, India; IISc Mathematics Initiative, Indian Institute of Science, Bangalore 560012, India; Department of Bioengineering, Indian Institute of Science, Bangalore 560012, India

**Keywords:** Membrane proteins, EMT, transcriptome, cancer, web server

## Abstract

Conventional approaches for analyzing ion channels in cancer primarily focus on the detection of membrane potential to record the polarization states of the cellular membrane. Although these approaches provide meaningful insights during validation, they face challenges in discovery, especially in identifying the ion channels that govern the membrane potential. Moreover, ion channels are known to exhibit context-dependent activity that often depends on the cancer type and metabolic state of the tumor. Here, we developed a computational framework, ONCOchannelome, that allows the identification of potential ion channels via transcriptomic datasets. We collected publicly available RNA-Seq datasets corresponding to 2421 normal, 5442 primary tumor and 588 metastatic samples of 15 tumor types. Using the data, we designed a computational strategy to merge the datasets, identifying differentially expressed ion channels in the tumor and metastatic states and aiding in the determination of ion channels in different tumor states. We subsequently developed strategies to determine the correlation of altered ion channels with epithelial-to-mesenchymal transition and to identify differentially coexpressed ion channel networks along with their possible transcription factors. The ONCOchannelome allows visualization of altered ion channels in the tumor and metastatic states in addition to revealing changes resulting in changes in biological processes and molecular functions as the tumor progresses. The ONCOchannelome enhances our understanding of the dependence of ion channels in different tumors in addition to revealing their observed alterations in progressing from the primary tumor state to the metastatic state.

## 1. Background

Ion channels, the sequentially and structurally distinct membrane proteins expressed in all cell types, are known for their ability to selectively permeabilize ion species within and across cell membranes. Furini and Domene reported that the human genome encodes over 400 distinct ion channels, accounting for approximately 2% of all identified genes (1). Traditional methods for studying these proteins lie within the scope of biophysicists and physiologists, who are primarily assessing physiological processes, including membrane excitability and epithelial transport. However, in recent years, increasing evidence has suggested that ion channels are involved in processes far beyond traditional physiological functions. Understanding the chemistry, electric potential and mechanics of ion channels could aid in regulating epigenetic and gene expression-related activities in a cell (2), (3), (4). Additionally, ion channels function as transcription factors and participate in cellular signaling through molecular interactions with macromolecular complexes (5), (6).

Aberrant expression of ion channels has been reported in multiple tumors (7), (8), (9), (10), (11). Essentially, the altered ion channel landscape or "channelome" in tumor cells appears to play an active role in processes such as neoplastic transformation, tumor progression, adaptation to the tumor microenvironment, metastasis, and therapy resistance (12), (13). Unlike oncogenes, ion channels are generally not considered to trigger the onset of cancer, but they are known to drive oncogenic processes by modulating factors such as calcium signaling, the membrane voltage, the sodium concentration, and the cellular pH (13). These observations, along with experimental interferences demonstrating associations of altered ion channels and tumor progression, resulted in the formation of the term “oncochannels” (13). In parallel, several pharmacological modulators of ion channels are already in clinical use or are in clinical trials (14). The overexpression of some oncochannels could qualify them as tumor-associated antigens suitable for vaccine development (15). Additionally, their presence at the cell surface would also allow for selective targeting. Specific antibodies with various strategies are already being used to exploit ion channels, demonstrating success in preclinical models and clinical trials (16), (17), (18).

Despite recent progress, characterizing ion channels continues to present theoretical and computational challenges. The involvement of ion channels in multiple physiological processes, along with their role in the development of cancer, increases the challenges in assigning specific mechanisms to ion channels in cancer hallmarks (19). Additionally, their small size, dynamic nature and diversity present difficulties in obtaining high-resolution structural information for understanding complex interactions with neighboring molecules (20), (21). Moreover, molecular dynamics simulations of ion channels face limitations due to computational constraints, including the accuracy of force fields, which lack the ability to capture biologically relevant timescales (22). The lack of annotations for many ion channels in cancer can be attributed to these limitations. To facilitate the detection of potential ion channels that could be targeted in various tumors, we developed the ONCOchannelome, a computational framework used to explore deregulated ion channels across tumors. Omics technologies such as transcriptomics aid in understanding various diseases, including stroke, diabetes, and cancer (23).

## 2. Methods

### 2.1 Databases used

For the functionality of the ONCOchannelome described herein, curated ion channel lists from the literature and gene symbols established in the Human Genome Organization Gene Nomenclature Committee (HGNC) database were used.

### 2.2 Input data

To develop the computational framework publicly available transcriptome profiling data accessible from The Cancer Genome Atlas (TGCA), Genotype Tissue Expression (GTEx) and Integrative Clinical Genomics of Metastatic Cancer (MET500) data resources were utilized. The cancer cohorts included breast invasive carcinoma (BRCA), colon adenocarcinoma (COAD), esophageal carcinoma (ESCA), head and neck squamous cell carcinoma (HNSC), pancreatic adenocarcinoma (PAAD), prostate adenocarcinoma (PRAD), skin cutaneous melanoma (SKCM), thyroid carcinoma (THCA), bladder urothelial carcinoma (BLCA), cholangiocarcinoma (CHOL), glioblastoma multiforme (GBM), kidney chromophobe (KICH), lung adenocarcinoma (LUAD), lung squamous cell carcinoma (LUSC), and stomach adenocarcinoma (STAD) (Figure 1).

**Figure 1:**
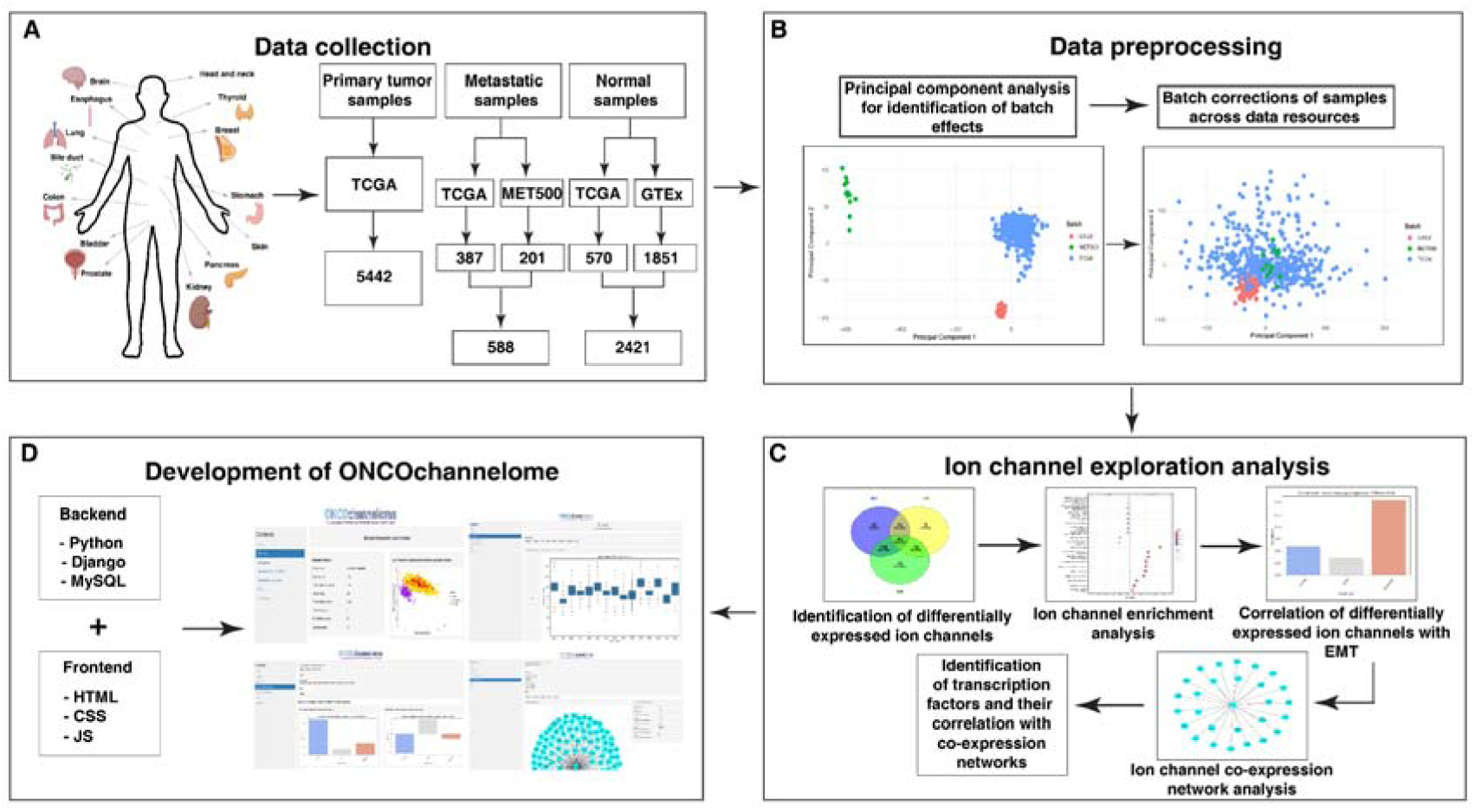
Depiction of the overall workflow used to develop the computational framework ‘ONCOchannelome’ to investigate altered ion channels across tumor types.

### 2.3 Preprocessing

Normal individual data from the GTEx were selected on the basis of age, sex and tumor sample ratio normalization. Principal component analysis (PCA) was performed across samples belonging to a specific tumor type, and batch effects were identified. The R Bioconductor limma (v.3.60.6) (24), afffy (v.1.82.0) (25) and sva (v.3.52.0) (26) packages were utilized to perform batch corrections on the basis of the sample group as well as the sample type specifically via the removeBatchEffect function from the limma package (S1 Fig and S2 Fig).

### 2.4 Detection of alterations in ion channels

**The** R Bioconductor limma package was used to estimate the differentially expressed genes (DEGs) via the Bayes method. DEGs were identified for 3 subgroups: tumor vs normal (TN), metastatic vs. tumor (MT) and metastatic vs. normal (MN). Any gene with an adjusted p value < 0. 05 values calculated via the Benjamini□Hochberg method of the false discovery rate and log2FC |0.6| (corresponding to FC > 1.5 or < 0.66) between the conditions were considered DEGs. The number of altered ion channels in each subgroup was given as a subset of the total DEGs obtained as described below.

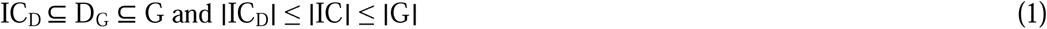

where G is the total number of transcripts, D_G_ is the total number of DEGs, IC is the total number of curated ion channels and IC_D_ is the total number of DE ion channels.

### 2.5 Gene Ontology enrichment

We performed Gene Ontology Biological Process, Molecular Function and Cellular Compartment gene set enrichment analysis (GSEA) on the IC_D_ in each subgroup via the R Bioconductor clusterProfiler library (v.4.2.2) (27). The minimum gene size was set to 5, the maximum gene size was set to 300, the p value cutoff was set to 0.05, and the number of permutations was set to 10000. We show the significantly enriched terms for the IC_D_ in each subgroup.

### 2.6 Construction of ion channel coexpression networks

To construct a state-specific ion channel network, we provided the transcriptome profiles corresponding to the IC_D_ to a weighted gene coexpression network analysis (WGCNA) library (v.1.73) (28), an R Bioconductor package. The expression profiles were preprocessed via the goodSampleGenes function, and the genes without any variance across samples were filtered out. A scale-free network model was generated, followed by construction of an adjacency correlation matrix to obtain an appropriate soft threshold power. The adjacency matrix was further transformed into a topological overlap matrix (TOM), resulting in a dissimilarity matrix. Thereafter, we performed hierarchical clustering via the DynamicTreeCut algorithm, and significant gene modules were selected. The expression profiles of each of the modules summarized in the form of module eigengenes were correlated with the binary trait values (normal, primary tumor, and metastatic). Gene significance (GS) and module membership (MM) values were calculated and further used to generate GS vs MM plots to estimate the ion channel associations with binary traits. The coexpressed gene modules were identified as those whose p value was <= 0.05. The soft threshold values interpreted via scale-free network topology fit and the minimum module size defined on the basis of the IC_D_s obtained for individual tumor types in each subgroup of identified ion channels are listed in Table S1.

### 2.7 Determination of the correlation of ion channel coexpression networks with epithelial to mesenchymal transition (EMT)

Single-sample gene set enrichment analysis (ssGSEA) was performed on the epithelial and mesenchymal gene lists from the transcriptome profiles of tumor types that contained metastatic samples. The gene lists were obtained from a previously published study (29). We quantified the enrichment of the gene signatures independently via gene set enrichment analysis in the Python (GSEAPY) library (30). The enrichment scores were further normalized for the gene sets and used to calculate their correlation with the DE ion channels via Spearman correlation.

### 2.8 Computation of the correlation of coexpressed network-specific transcription factors with p values

The coexpressed IC_D_ modules were analyzed via Enrichr (31), and we determined the transcription factors that might control the ion channel coexpressed networks. The gene set libraries from Enrichr used were as follows: ChIP enrichment analysis (ChEA), Enrichr for protein□protein interactions (PPIs), Encyclopedia of DNA elements (ENCODE) transcription factor (TF) ChIP-seq 2015, TRANScription FACtor database (TRANSFAC) and JASPAR position weight matrix (PWMs), ENCODE and ChEA consensus TFs from ChIP-X, and transcriptional regulatory relationships unraveled by sentence-based text mining (TRRUST). The transcription factors with a p value cutoff of 0.05 and with evidence of presence in humans were shortlisted. The analysis was updated last on 29^th^ March 2025. Thereafter, Pearson correlation along with t-distribution-based p values were calculated between the expression values of the transcription factors, and WGCNA was used to obtain module eigen genes to associate the coexpressed gene modules with the determined transcription factors.

### 2.9 ONCOchannelome web implementation

The ONCOchannelome was formulated as a web interface developed via Python (v.3.10.11) scripts based on the Django (v.5.0.3) framework. The data generated through the analysis were stored in the MySQL relational database management system and can be accessed by the users through the web interface via a link (https://www.oncochannelome.co.in/). The front-end design of the web interface was accomplished via hypertext markup language (HTML), bootstrapping, cascading style sheets (CSSs) and a Java script (JS).

## 3. Results

### 3.1 ONCOchannelome modules

The ONCOchannelome comprises four modules: Browse by tumor type, ion channel retrieval, EMT correlation and the ion channel coexpression network (Figure 2).

**Figure 2:**
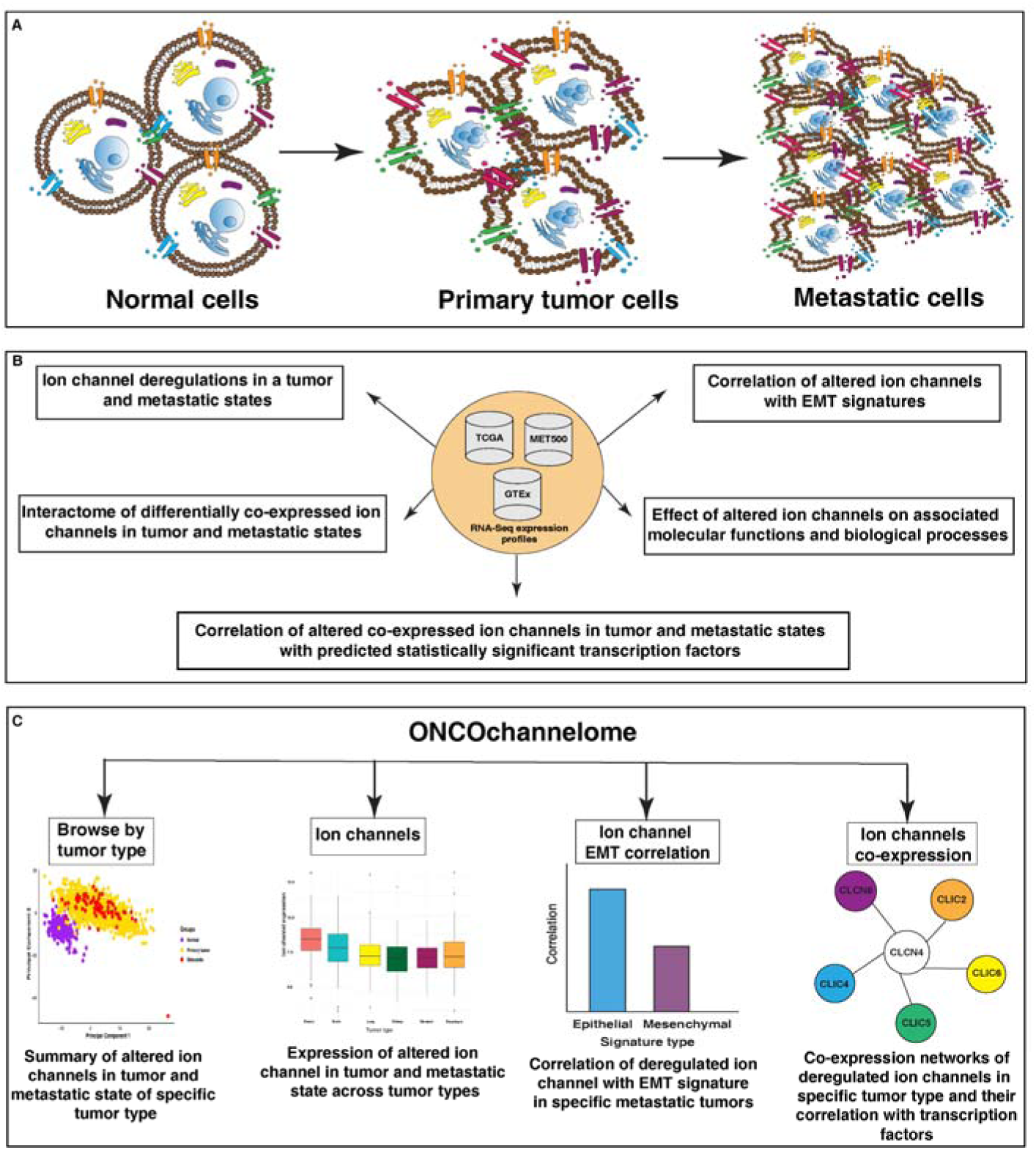
Schematic overview of the ONCOchannelome: A) The transformation between normal to tumor and tumor to metastatic states during tumor progression involves the cells to undergo molecular alterations resulting in significant changes in the cell morphology that reflect the traits required for metastatic invasion. B) The ONCOchannelome consists of modules that facilitate the determination of differentially expressed (DE) ion channels in the tumor and metastatic states of 15 tumor types, correlate the DE ion channels with epithelial and mesenchymal gene signatures, construct ion channel coexpression networks, provide details on the biological processes and molecular functions of DE ion channels and provide correlations with potential transcription factors. C) Schematic of the ONCOchannelome. The computational framework allows users to explore ion channels via four modules: Browse by tumor type, ion channels, ion channel-EMT correlation and ion channel coexpression.

The browse by tumor type module allows users to explore the tumor types in terms of the overall deregulations observed in the ion channels. The ion channel retrieval module allows searches of the expression and transcriptomic changes of a specific ion channel across tumor types. The ion channel EMT correlation module can be used to determine the correlations between altered ion channels in a tumor type in a tumor state and epithelial and mesenchymal gene signatures. The fourth module, ion channel coexpression, can be used to determine the ion channels that are coexpressed with the ion channel of interest in a tumor type in a tumor state. The significant associations of the ion channel coexpression network with transcription factors can also be identified through this module.

### 3.2 Browse by tumor type

We included 5442 primary tumor samples; 588 metastatic samples and 2421 normal samples distributed across 15 tumor types from 14 primary sites: BRCA, COAD, ESCA, HNSC, PAAD, PRAD, SKCM, THCA, BLCA, CHOL, GBM, KICH, LUAD, LUSC and STAD (Figure 3A).

**Figure 3:**
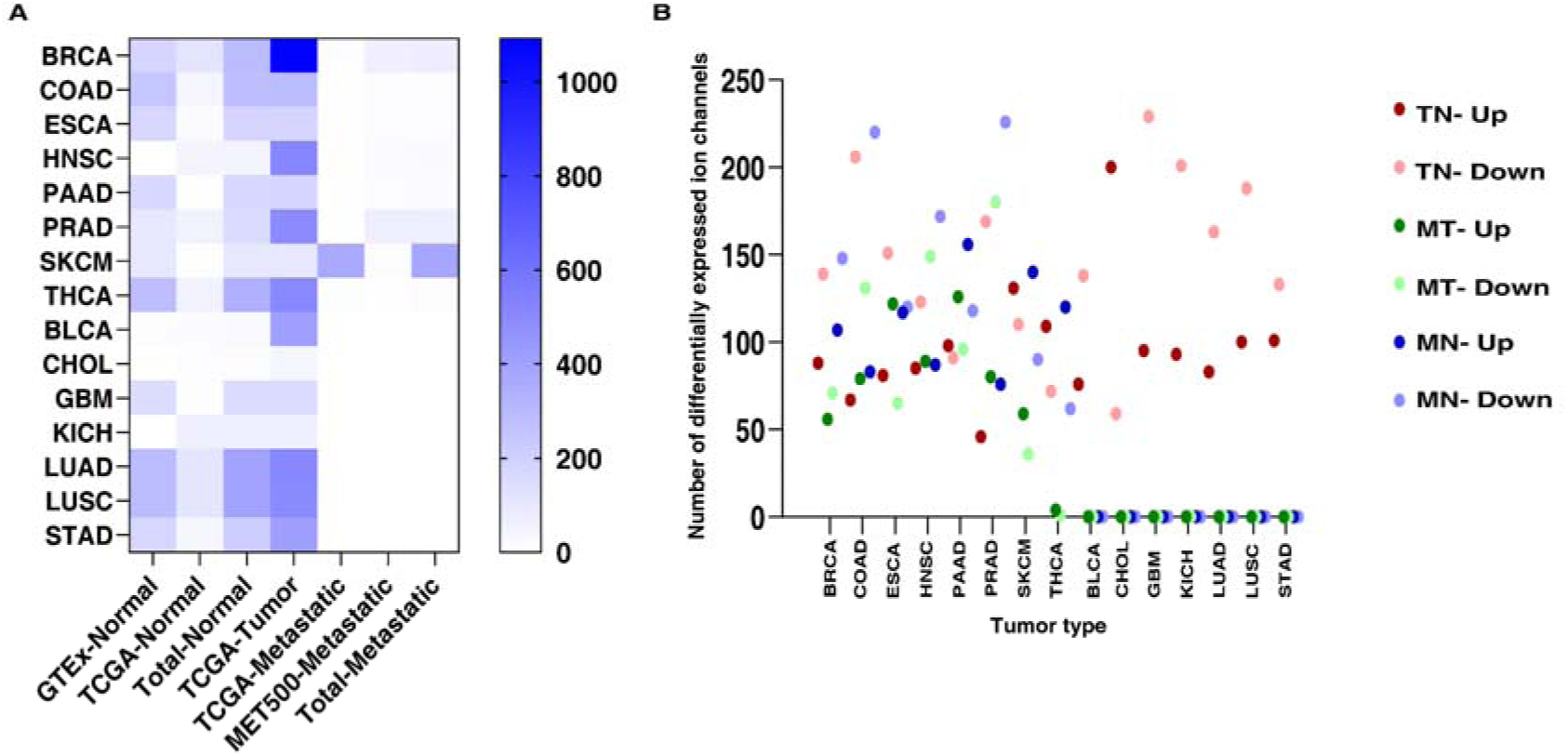
Summary of samples collected and differentially expressed ion channels across tumor types: A) Number of samples available across tumor types across normal, primary tumor and metastatic states. B) Number of upregulated and downregulated ion channels across tumor types across primary tumor and metastatic states.

Previously, we reported DE ion channels in the tumor and metastatic states of breast cancer (9). Ion channels were identified as potential candidates for determining the prognosis of patients with breast cancer. Given the nonuniform expression pattern of ion channels in a specific sample state, the ion channels overexpressed in a particular cancer type may or may not overlap with the ion channels overexpressed in other cancer types or sample states. Furthermore, the details available in the literature are scattered and not easily accessible. In view of this, we collected a list of ion channels and overlaid them to represent the unique patterns of altered expression observed in each tumor type and tumor state (Figure 3B).

The impact on the molecular functions, biological processes and cellular compartments due to alterations observed could also be established from the datasets analyzed.

### 3.3 Ion channel retrieval

The module includes details on 493 ion channels. For each ion channel, information retrieved from public data resources such as FireBrowse, TumorPortal, GTEx, DepMap, PubMed, Harmonizome, Cancerscem and MET500 was linked with ONCOchannelome. Thereafter, the expression levels after performing batch correction to combine the datasets from the different data sources were plotted to depict the expression profiles for each individual ion channel across tumors. ONCOchannelome-enabled altered ion channel visualization will help users identify the extent of deregulation of a particular ion channel across tumors in different tumor states.

### 3.4 Ion channel EMT correlation

Previously, we reported the correlation of altered ion channels with EMT gene signature scores, indicating mesenchymal phenotypes in patients with breast cancer and esophageal squamous cell carcinoma (9), (8). EMT signature scores are calculated via previously published signature gene sets. ssGSEA enrichment scores for epithelial and mesenchymal signature gene sets were computed across transcriptome profiles of the available metastatic tumor samples. Normalized enrichment scores revealed distinct patterns of epithelial and mesenchymal signatures across tumor types. Thereafter, the correlation of the DE ion channels in a specific tumor type with the calculated EMT scores was computed. Through the ONCOchannelome ion channel EMT correlation panel, users can select an ion channel of interest and the tumor type and submit a query to evaluate the relationship between epithelial and mesenchymal gene signatures and the ion channel in normal, tumor and metastatic states of a particular tumor type.

### 3.5 Ion channel coexpression

The association between altered ion channels and strongly coexpressed ion channels, along with their extent of deregulation, can facilitate the identification of possible relationships between ion channels on the basis of their expression patterns across different samples. ONCOchannelome allows users to explore this association of altered ion channel expression with the expression of coexpressed ion channels. Furthermore, it allows users to identify the transcription factors that may regulate the expression of the coexpressed network. This information might provide insights into the regulatory mechanisms underlying the observed ion channel expression patterns in a particular tumor type.

### 3.6 Case study

#### 3.6.1 Comparative analysis of ion channel modulation in histological subtypes of lung cancer

ONCOchannelome can be utilized to compare the alterations in ion channels observed in adenocarcinoma versus squamous cell carcinoma. We performed a comprehensive comparison study between LUAD and LUSC. The ONCOchannelome consisted of 397 normal samples from both LUAD and LUSC patients, whereas it consisted of 513 primary tumor samples from LUAD patients and 498 primary tumor samples from LUSC patients. Eighty-three ion channels were upregulated in LUAD, and 100 ion channels were upregulated in LUSC. Similarly, approximately 163 and 188 ion channels were downregulated in the two tumor types, respectively. Overlaying the altered ion channels in the two subtypes indicated an overlap of 208 ion channels. Alternatively, 38 and 80 ion channels were identified as subtype-exclusive ion channels in LUAD and LUSC, respectively. The ion channels *GJB7*, *KCTD15*, *SLC10A6*, *CACNA2D3*, *KCNK5*, *CRACR2B* and *SLC41A2* were found to have divergent expression patterns in the LUAD and LUSC subtypes. The variability observed in terms of alterations was reflected in the enriched biological processes and molecular functions as well. Two biological processes, glycerol transmembrane transport and inhibitory synapse assembly, were identified as being exclusively activated in the LUSC subtype. Similarly, the biological processes inorganic cation transmembrane transport, plasma membrane organization and plasma membrane phospholipid scrambling were activated only in the LUAD subtype. The molecular functions carbohydrate transmembrane transporter activity, polyol transmembrane transporter activity and glycerol transmembrane transporter activity were exclusively activated, and symporter activity, active monoatomic ion transmembrane transporter activity, antiporter activity, organic acid transmembrane transporter activity and carboxylic acid transmembrane transporter activity were exclusively suppressed in the LUSC subtype. Similarly, intracellular calcium-activated chloride channel activity, GABA-gated chloride ion channel activity, high-voltage gated calcium channel activity and GABA receptor activity were activated, whereas monoatomic cation transmembrane transporter activity, metal ion transmembrane transporter activity, inorganic cation transmembrane transporter activity, calcium channel activity and calcium ion transmembrane transporter activity were suppressed exclusively in the LUAD subtype. The cellular compartments observed to be activated in LUAD involved regions specific to neuronal transport, indicating its potential in brain metastasis, whereas the ion channels involved in neuronal mechanisms were suppressed in the LUSC subtype. Coexpression networks of ion channels showing divergent expression patterns provided further insights into their importance in terms of tumorigenicity. For example, *GJB7* was shown to be elevated in LUSC and suppressed in LUAD. The coexpression network of ion channels in LUSC revealed that *GJB7* was highly significant within the network (0.96, p value: 1.20e^--52^), whereas the network was negatively correlated with normal samples (-0.89, p value: 2.00e^--300^) and positively correlated with tumor samples (0.89, p value: 2.00e^--300^). All the ion channels in the network were downregulated. The transcription factor *EZH2* was highly positively correlated with the coexpression network (0.87, p value: 2.28e^-238^), and *E2F1* was moderately positively correlated with the network (0.67, p value: 6.44e^-122^). In LUAD, the coexpression network indicated that *GJB7* was highly significant within the network (0.87, p value: 1.30e^-51^), with the network being positively correlated with normal samples (0.8, p value: 2.00e^-207^) and negatively correlated with tumor samples (-0.8, p value: 2.00e^-207^). All the ion channels, excluding *AQP6*, a water transporter, were downregulated in the network. *JUN* was identified as the transcription factor (p value: 0.03) regulating *GJB7* downregulation along with the ion channels *SLC10A4*, *GABRB3*, *SLC10A6*, *SLC30A8*, *KCNIP1*, *KCNJ12*, *KCNC4*, *SLC16A12*, *KCNAB1*, *KCNA5*, *GJC1*, *GRIN2A*, *GJA5*, *KCNMB1*, *CACNG4* and *KCNJ3* (Figure 4,5).

**Figure 4:**
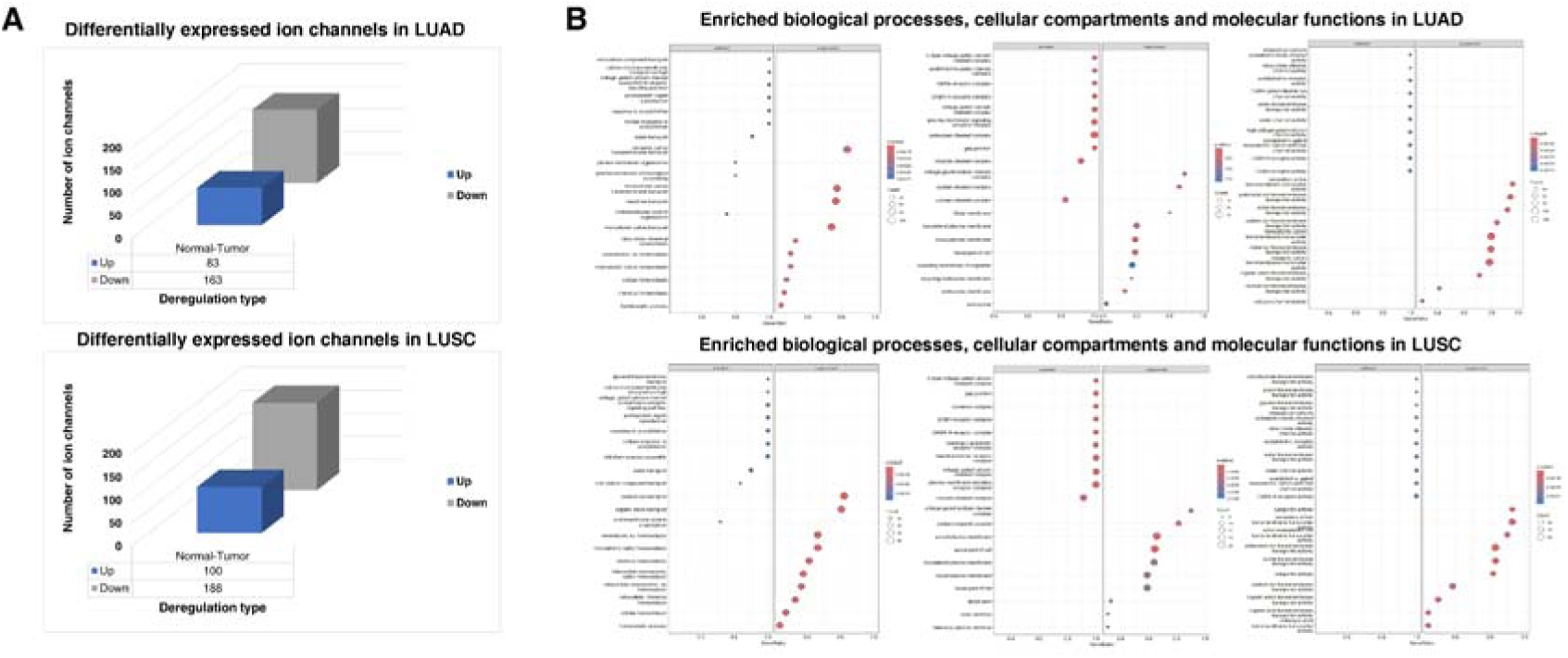
Altered ion channels in histological subtypes of lung cancer. A) Number of differentially expressed ion channels in LUAD and LUSC samples. B) Dot plot of activated and suppressed biological processes, molecular functions and cellular compartments corresponding to differentially expressed ion channels in LUAD and LUSC.

**Figure 5:**
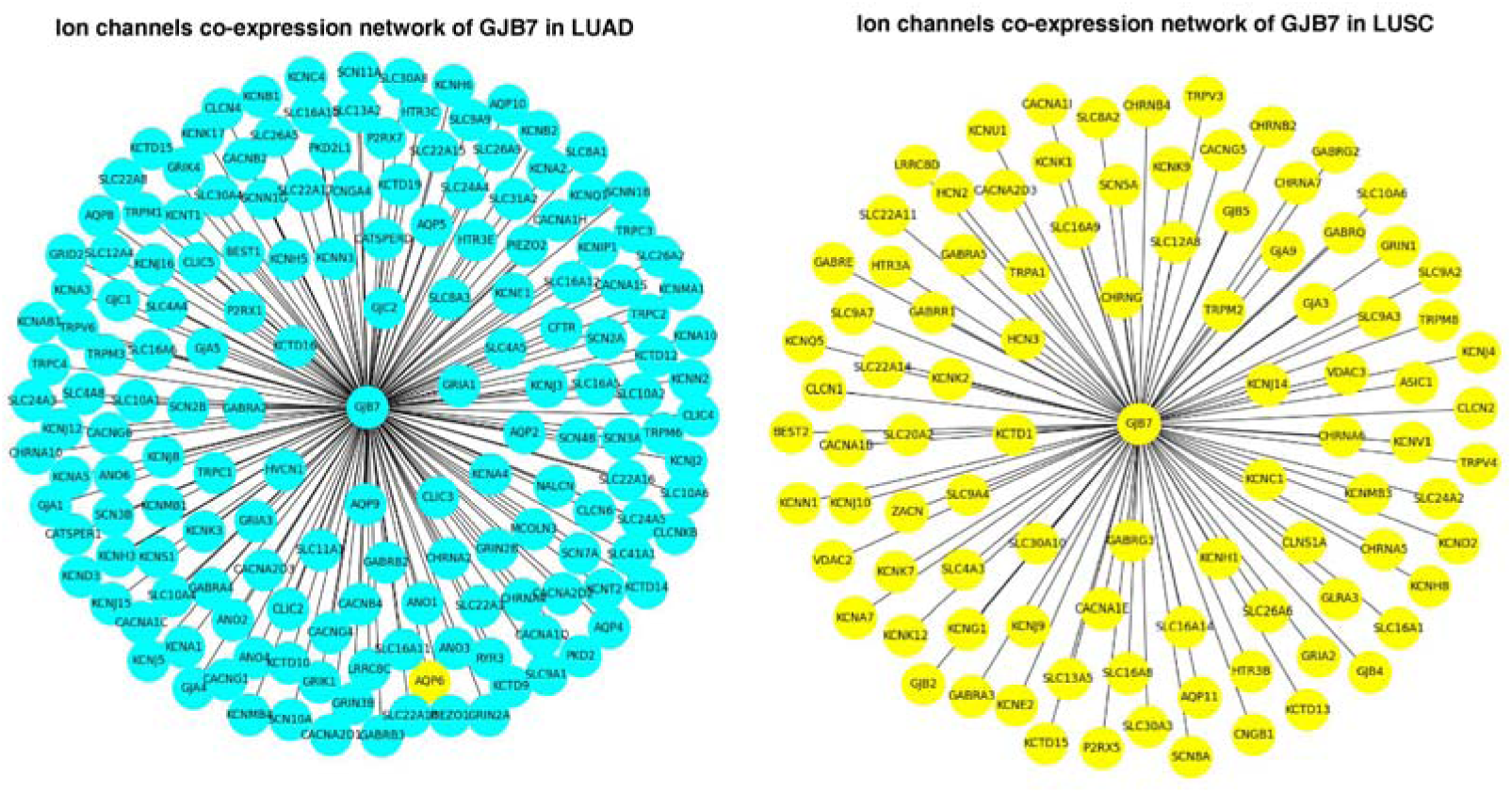
Ion channel coexpression network of ion channels coexpressed with GJB7 in LUAD and LUSC.

#### 3.6.2 Alterations in ion channels in the tumor and metastatic states of skin cutaneous melanoma

The ONCOchannelome has eight tumors with samples corresponding to the primary tumor and metastatic states. We report a comprehensive overview of the ion channels found to be altered in the tumor and metastatic states of SKCM. The ONCOchannelome consisted of 102 normal and primary tumor samples corresponding to SKCM and 378 metastatic samples. Differential gene expression analysis was carried out to compare TN, MT and MN, which revealed 43 DE ion channels that overlapped across the states. 27, 5 and 31 ion channels were exclusively DE in TN, MT and MN, respectively. Among the 43 ion channels, 10 exhibited uniform expression patterns: either consistently upregulated or downregulated across all SKCM states. The remaining 33 ion channels demonstrated consistent deregulation when normal tissue was compared with both tumor and metastatic samples. However, when tumor samples were compared with metastatic samples, the expression of these ion channels was reversed. The gene set enrichment analysis obtained from ONCOchannelome revealed a unique set of enriched molecular functions in each comparison state, with only the function of ‘identical protein binding’ being common to normal samples compared with tumor and metastatic samples. Similarly, the molecular functions adenyl nucleotide binding, adenyl ribonucleotide binding and neurotransmitter receptor activity overlapped between the metastatic samples and the tumor and normal samples. Among the ten ion channels, *GJB2*, *GLRB*, *GRIA3*, *GRID2*, *HTR3A*, *KCND2*, *KCTD16*, *PIEZO2* and *SLC26A4* were positively correlated with mesenchymal gene signatures. Among the ion channels identified as exclusively altered in tumors compared with the metastatic state, *CACNA1E*, *SCN7A*, *CACNA1G* and *KCNK1* were more positively correlated with mesenchymal gene signatures than with epithelial gene signatures, indicating their significance in tumor progression (Figure 6).

**Figure 6:**
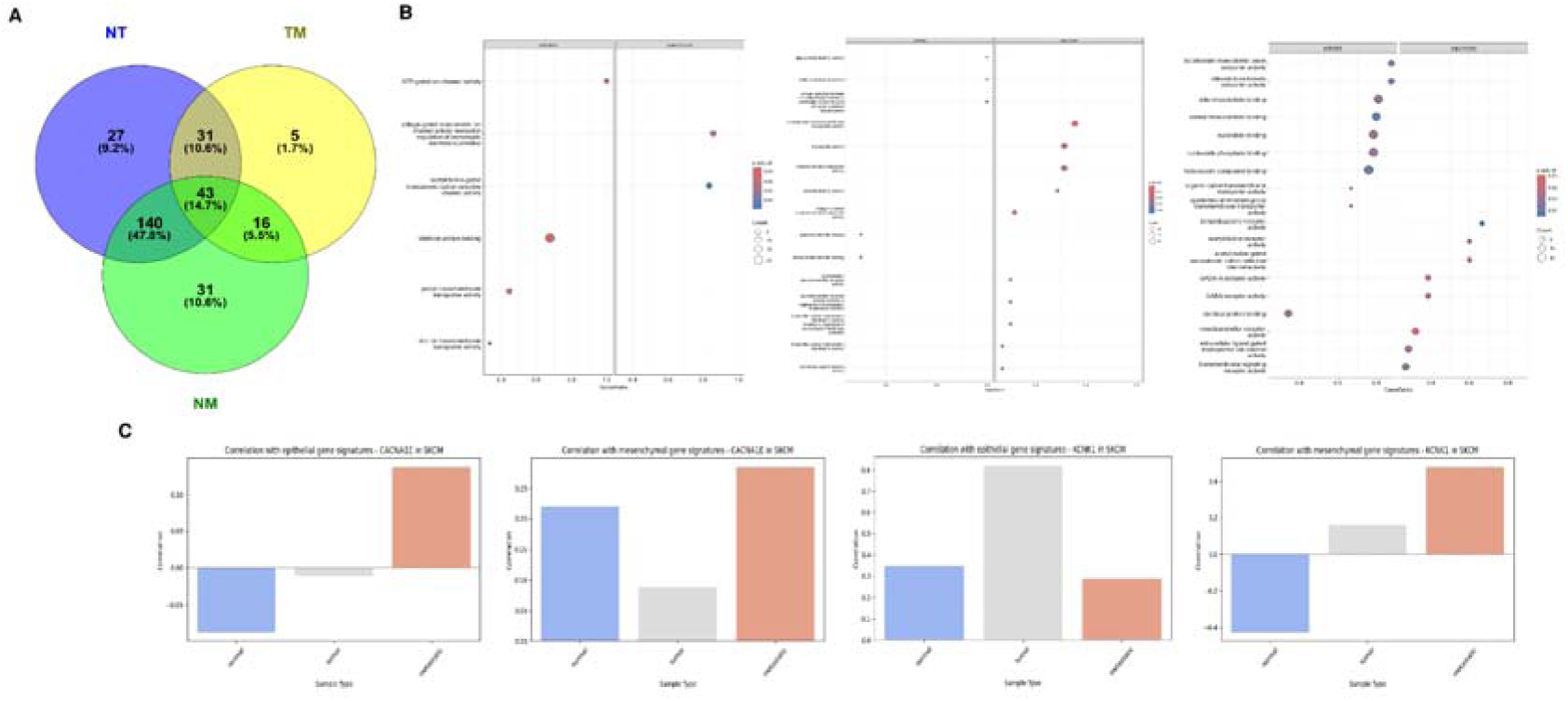
Ion channels altered in the tumor and metastatic states of skin cutaneous melanoma: A) Overlaps in differentially expressed (DE) ion channels in tumor samples compared with normal samples (TN), metastatic samples compared with tumor samples (MT) and metastatic samples compared with normal samples (MN). B) Dot plot of activated and suppressed molecular functions corresponding to DE ion channels in the TN, MT and MN states. C) Correlations of epithelial and mesenchymal gene signatures with *CACNA1E* and *KCNK1* in normal, tumor and metastatic expression profiles.

### 3.7 User interface and information retrieval

ONCOchannelome has a simple user interface for data exploration of altered ion channels across tumor types. The interface enables users to identify alterations in ion channels and retrieve their significance in various biological processes and molecular functions. Hyperlinks are provided to various projects that allow users to correlate altered ion channels with other studies that have identified mutations and expression patterns in ion channels in various tumors via different datasets. Correlation with EMT gene signatures can be obtained for individual ion channels altered in specific tumors. Coexpression of ion channels with other significantly altered ion channels would allow the user to look for gene clusters that may function together, resulting in downstream processes. Potential transcription factors can also be identified to hypothesize that the regulation of biological processes is deregulated within the tumor, which may provide insights into the regulatory mechanisms underlying the observed gene expression patterns.

## 4. Discussion

The ONCOchannelome computational framework was designed to devise a computational method to identify oncochannels across tumor types and to enhance our understanding of ion channels in tumor progression. It represents a computational platform to explore ion channel deregulation along with providing details in terms of their correlation with EMT. ONCOchannelome goes beyond currently available resources, namely, IUPHAR-DB (32), ChanFAD (33) and Channelpedia (34), with a focus on ion channels and their involvement in cancer. The ONCOchannelome can be used for (i) exploration of alterations in ion channels in tumor and metastatic states across tumor types; (ii) correlation of specific ion channels identified to be altered in particular tumors with epithelial and mesenchymal gene signatures; (iii) retrieval of biological processes, molecular functions and cellular compartments with the possibility of being activated or suppressed in the deregulation of specific ion channels; and (iv) prediction of potential transcription factors with the potential to regulate altered ion channels. (v) determination of the ion channel interactome on the basis of statistical significance from transcriptome gene coexpression analysis and visualization of results.

Currently, ONCOchannelome incorporates processed data via the developed computational pipeline using the TCGA, MET500 and GTEx datasets, which consists of 5442 primary tumor samples, 588 metastatic samples and 2421 normal samples with information on 493 ion channels. Hyperlinks toward high-throughput data analysis projects, such as FireBrowse, TumorPortal, GTEx, PubMed, Harmonizome, Cancerscem and MET500, which allows broad-spectrum comparisons of ion channels across datasets other than those included in the computational framework, are also included. Notably, altered ion channels across tumor types were detected. Furthermore, overlaps of altered ion channels in tumor and metastatic states of tumors were depicted. The distributions of RNA abundance in normal, tumor and metastatic states across tumors for each ion channel are provided in the form of boxplots. The correlations between EMT gene signatures and DE ion channels were assessed across normal, tumor, and metastatic states across tumor types. This correlation of altered ion channels in each tumor type with epithelial and mesenchymal gene signatures can be visualized in the form of bar plots. The plots could be utilized to observe associations that may reflect general EMT-related transcription shifts across heterogeneous samples of a specific tumor. Although the correlation analysis did not consider alternative variables such as histological subtype, tumor stage or demographics, the observed correlations were consistent with findings from previous studies. For example, the mRNA expression of transient receptor potential cation channel subfamily C member 1 (*TRPC1*) was reported to be strongly associated with the mesenchymal phenotype in endometrial carcinoma (35). Our methodology revealed that the expression of *TRPC1* in BRCA, COAD, HNSC, PRAD and SKCM was more strongly correlated with mesenchymal gene signatures than with epithelial gene signatures, which may indicate an aggressive tumor phenotype in these tumors. Similarly, potassium voltage-gated channel subfamily Q member 1 (*KCNQ1*) was identified to have potential significance in epithelial cancers*, in which low KCNQ1 expression* was reported to promote colon adenocarcinoma (36). The expression of *KCNQ1* was associated with epithelial cell plasticity, indicating that it promotes epithelial features (36). Our methodology revealed that the expression of *KCNQ1* in multiple tumors, including COAD, ESCA, PAAD, PRAD and THCA, was more strongly correlated with epithelial gene signatures than with mesenchymal gene signatures. Similarly, gap junction *GJB7* has been previously reported to harbor somatic mutations in gastric and colorectal cancers (37). Our study revealed transcriptional alterations in *GJB7* in lung carcinoma. Furthermore, gene coexpression networks, along with indications of the upregulation and downregulation of each ion channel in the network, are provided. The correlation values of the network with binary traits, i.e., normal, tumor and metastatic traits, further establish the statistical relevance of the network. The computational framework also includes transcription factor identification on the basis of the gene coexpression networks generated and their correlation with the complete network. Furthermore, we provide two case studies to help users explore the framework for their research.

### Limitations

The utility of the computational framework ONCOchannelome is best realized during the discovery phase, as it facilitates the generation of hypotheses by identifying potential ion channels altered in specific tumor types. Translational interpretation of the computationally established ion channels would require further experimental validation. Furthermore, the imbalance in the sample distribution across normal, tumor and metastatic states may constrain the statistical power for detecting subtle alterations specific to progression. Nonetheless, the utilized dataset provides a robust and valuable foundation for comparative analysis, and the devised framework could be used for larger datasets on availability. The ONCOchannelome provides a broader view of the association of EMT with DE ion channels. The incorporation of clinical metadata may be needed to enhance and validate context-specific EMT dynamics. The transcription factors predicted for DE ion channels derived from multiple curated databases are computational in nature. These findings offer valuable insights and serve as the basis for further investigations.

## 5. Conclusion

In summary, the ONCOchannelome serves as a comprehensive resource for identifying tumor- and metastatic-related ion channels in 15 tumor types, exploring EMT-related correlations for each altered ion channel, and analyzing the relevance of deregulation of the ion channel in biological processes and molecular functions. The interface and the structure of the framework allow for the inclusion of additional cancer types on the basis of the availability of datasets. Currently, ONCOchannelome lacks the inclusion of other omics layers in terms of ion channel alteration data, but it could be explored with the availability of new datasets.

## 6. List of abbreviations

HGNC: Human Genome Organization Gene Nomenclature Committee
TCGA: The Cancer Genome Atlas
GTEx: Genotype Tissue Expression
MET500: Integrative Clinical Genomics of Metastatic Cancer
BRCA: Breast invasive carcinoma
COAD: Colon adenocarcinoma
ESCA: Esophageal carcinoma
HNSC: Head and neck squamous cell carcinoma
PAAD: Pancreatic adenocarcinoma
PRAD: Prostate adenocarcinoma
SKCM: Skin cutaneous melanoma
THCA: Thyroid carcinoma
BLCA: Bladder urothelial carcinoma
CHOL: Cholangiocarcinoma
GBM: Glioblastoma multiforme
KICH: Kidney chromophobe
LUAD: Lung adenocarcinoma
LUSC: Lung squamous cell carcinoma
STAD: Stomach adenocarcinoma
EMT: Epithelial to mesenchymal transition
TN: Tumor vs normal
MT: Metastatic vs. tumor
MN: Metastatic vs. normal
GSEA: Gene set enrichment analysis
WGCNA: Weighted gene coexpression network analysis
TOM: Topological overlap matrix
GS: Gene significance
MM: Module membership
ssGSEA: Single-sample gene set enrichment analysis
GSEAPY: Gene set enrichment analysis in the Python
ChEA: ChIP enrichment analysis
PPI: Protein[protein interactions
ENCODE: Encyclopedia of DNA elements
TF: Transcription factor
TRANSFAC: TRANScription FACtor database
PWM: Position weight matrix
TRRUST: Transcriptional regulatory relationships unraveled by sentence-based text mining
HMTL: Hypertext markup language
CSS: cascading style sheets
PCA: Principal component analysis
DEG: differentially expressed genes

## 7. Declarations

### · Ethics approval and consent to participate

Not applicable

### · Consent for publication

Not applicable

### · Availability of data and materials

The ONCOchannelome is available at ‘https://www.oncochannelome.co.in/’.

### · Competing interests

The authors declare that they have no competing interests

### · Funding

The author(s) received no specific funding for this work.

### · Authors’ contributions

K.T.S.P.: Writing – review & editing, Writing – original draft, Visualization, Software, Methodology, Investigation, Formal analysis, Data curation. S.R.: Methodology, formal analysis. M.K.J.: Formal analysis, Writing – review and editing. J.S.: Writing – review & editing, Methodology, Supervision, Resources, Project administration, Investigation, Conceptualization.

## Acknowledgements

We would like to thank the authors of the manuscripts for making the datasets used in this study publicly available. We thank Harsh Pawar for helpful discussions and critical review. J.S. would like to thank the Council of Scientific and Industrial Research (CSIR), the Government of India [37WS(0114)/2023-24/EMR-II/ASPIRE] and the Indian Council of Medical Research (ICMR), Government of India, for research support. K.T.S.P. was supported by ICMR, Government of India [BMI/12(95)2021]. J.S. was a recipient of the Bio-CARe Women Scientists award from the Department of Biotechnology (DBT), Government of India.

## 9. Additional files

**Table S1:** Parameters used in WGCNA of ion channels in TNs, MTs and MNs across tumors

**S1 Fig:** PCA plot to identify batch effects in tumor types with normal, tumor and metastatic samples available

**S2 Fig:** PCA plot to identify batch effects in tumor types with normal and tumor samples available

## 10. Data availability statement

The ONCOchannelome is available at ‘https://www.oncochannelome.co.in/’.

## Notes

https://www.oncochannelome.co.in/

### Competing Interest Statement

The authors have declared no competing interest.

### Summary of Updates

Abstract was revised. Tables were removed and new Figure 1 was added. The manuscript has been restructured.

