## Supplementary material for "ONCOchannelome: A computational framework for investigating altered ion channels across tumor types": Table S1, S1 Fig, S2 Fig

**Supplementary data**

1. Supplementary figures

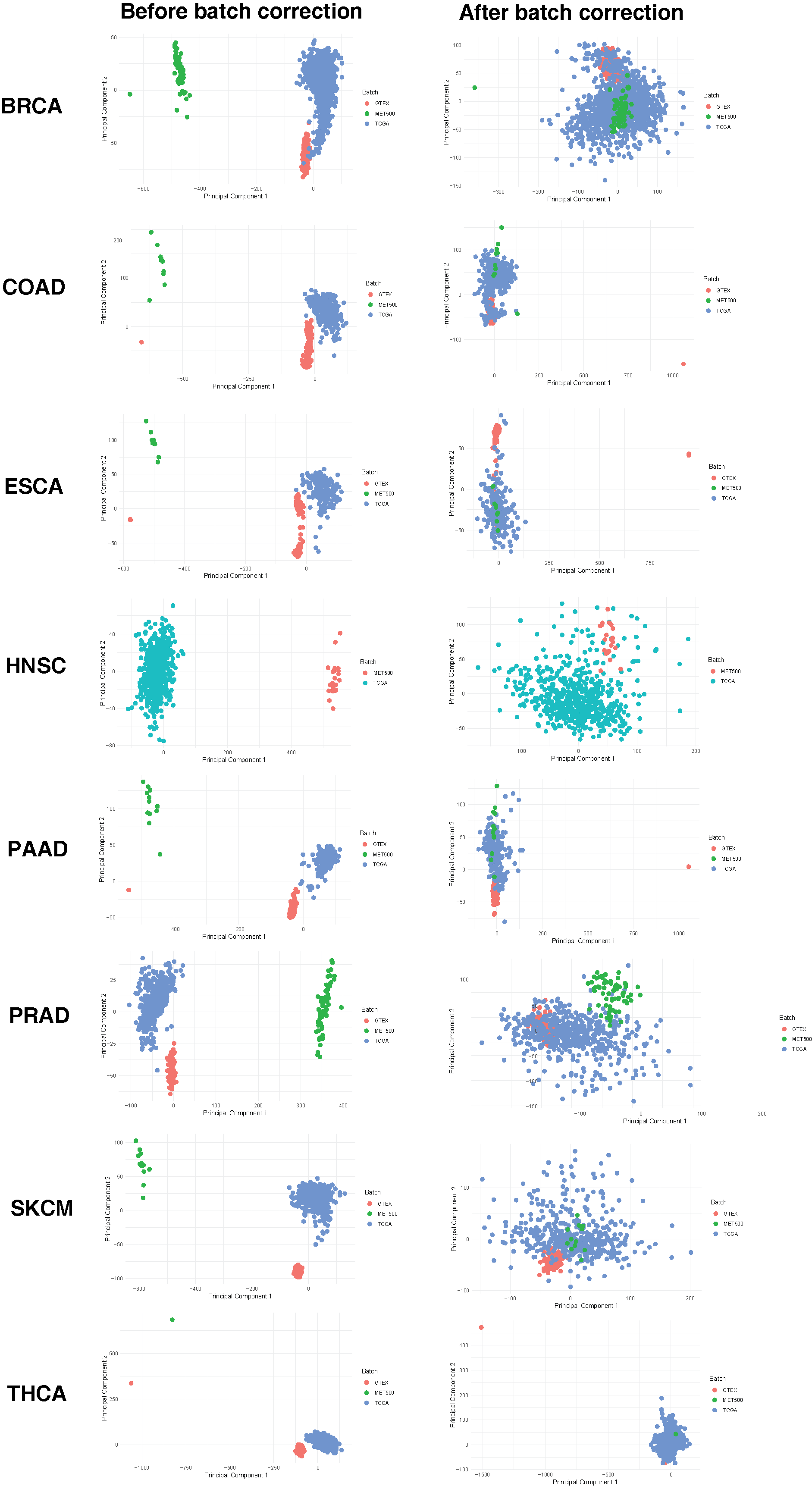

**S1 Fig:** PCA plot to identify batch effects in tumor types with normal, tumor and metastatic samples available

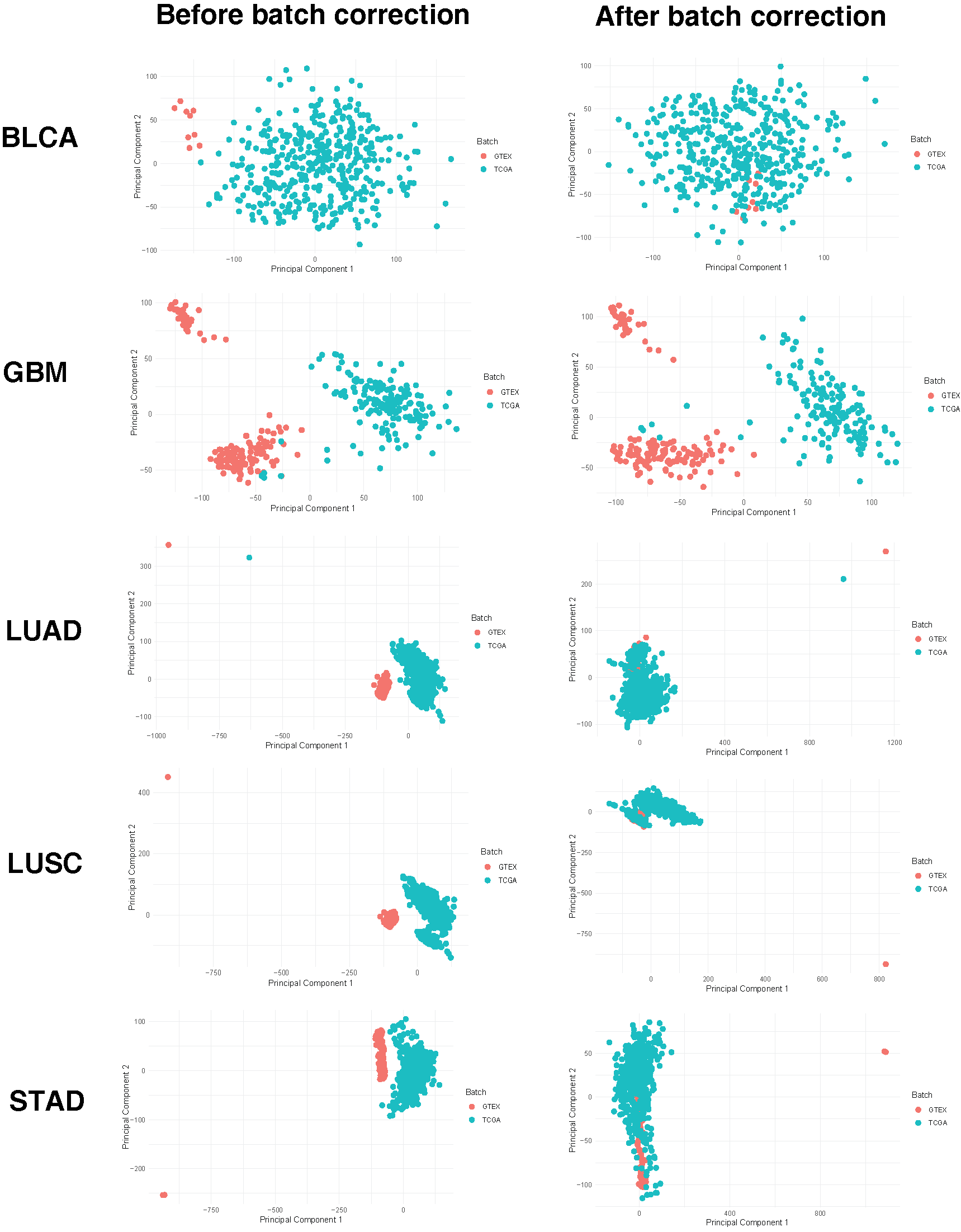

**S2 Fig:** PCA plot to identify batch effects in tumor types with normal and tumor samples available

**Supplementary tables**

**S1 table:** Parameters used in WGCNA analysis of ion channels in TN, MT and MN across tumors

| **Tumor type** | **Subgroup** | **Soft threshold power** | **Minimum module size** | **Deep split** | **Obtained number of modules** | **Chosen module** |
| --- | --- | --- | --- | --- | --- | --- |
| BLCA | TN | 12 | 30 | 4 | 3 | Turquoise |
| CHOL | TN | 18 | 30 | 4 | 3 | Blue, turquoise |
| GBM | TN | 14 | 30 | 4 | 4 | Blue, brown, turquoise |
| KICH | TN | 8 | 30 | 4 | 3 | Blue, turquoise |
| LUAD | TN | 10 | 30 | 4 | 3 | Blue, turquoise |
| LUSC | TN | 14 | 30 | 4 | 3 | Blue, turquoise |
| STAD | TN | 8 | 30 | 4 | 3 | Blue, turquoise |
| COAD | MN | 20 | 30 | 4 | 4 | Brown |
|  | MT | 8 | 30 | 4 | 3 | - |
|  | TN | 20 | 30 | 4 | 3 | Blue, turquoise |
| ESCA | MN | 9 | 30 | 4 | 4 | Blue, brown, turquoise |
|  | MT | 12 | 30 | 4 | 3 | Turquoise |
|  | TN | 12 | 30 | 4 | 4 | Blue, turquoise |
| HNSC | MN | 16 | 30 | 4 | 3 | Blue, turquoise |
|  | MT | 12 | 30 | 4 | 4 | Blue, brown, turquoise |
|  | TN | 10 | 30 | 4 | 4 | Blue, turquoise |
| PAAD | MN | 9 | 30 | 4 | 3 | Blue |
|  | MT | 14 | 30 | 4 | 4 | Blue |
|  | TN | 12 | 30 | 4 | 3 | Blue, turquoise |
| PRAD | MN | 7 | 30 | 4 | 3 | Blue, turquoise |
|  | MT | 7 | 30 | 4 | 3 | Turquoise, blue |
|  | TN | 12 | 30 | 4 | 3 | Blue, turquoise |
| SKCM | MN | 10 | 30 | 4 | 3 | Blue, turquoise |
|  | MT | 12 | 30 | 4 | 3 | Blue, turquoise |
|  | TN | 16 | 30 | 4 | 3 | Blue, turquoise |
| THCA | MN | 10 | 30 | 4 | 3 | Blue |
|  | MT | - | - | - | - | - |
|  | TN | 10 | 30 | 4 | 3 | Blue, turquoise |
| BRCA | MN | 16 | 30 | 4 | 4 | Blue, brown, turquoise |
|  | MT | 18 | 10 | 3 | 3 | Turquoise |
|  | TN | 10 | 30 | 4 | 3 | Blue, turquoise |
